# Inducible genome-wide mutagenesis for improvement of pDNA production by *E. coli*

**DOI:** 10.1101/2025.06.13.659583

**Authors:** Zidan Li, George Sun, Ibrahim Al’Abri, Yihui Zhou, Nathan Crook

**Affiliations:** Department of Chemical and Biomolecular Engineering, North Carolina State University, Raleigh, NC, USA; Bioinformatics Research Center, North Carolina State University, Raleigh, NC, USA

**Keywords:** plasmid DNA, *Escherichia coli*, strain engineering, genome mutagenesis

## Abstract

Plasmid DNA (pDNA) is a cost-driving reagent for the production of gene therapies and DNA vaccines. Improving pDNA production in the most common production host (*E. coli*) has faced obstacles arising from the complex network of genes responsible for pDNA synthesis, with the specific enzyme(s) limiting pDNA yield remaining unidentified. To address this challenge, we employed an inducible genome-wide mutagenesis strategy, combined with fluorescent screening, to isolate *E. coli* NEB 5α strains with enhanced pDNA production. Following selection, we successfully isolated an *E. coli* strain (M3) with elevated plasmid copy numbers (PCNs) across multiple origin types. Specifically, we observed a 5.93-fold increase in PCN for the GFP reporter plasmid, a 1.93-fold increase for the gWiz DNA vaccine plasmid, and an 8.7-fold increase for the pAAV-CAGG-eGFP plasmid, all of which contain pUC origins. In addition, plasmids with p15A and pSC101 origins showed 1.44-fold and 1.68-fold increases in PCN, respectively. Whole-genome sequencing of the adapted strain M3 identified 85 mutations, including one in *recG*, which encodes an ATP-dependent DNA helicase. Replacement of the mutant *recG* with its wild-type counterpart in the mutant strain resulted in a 63% reduction in PCN, but the *recG* mutation alone was insufficient to increase PCN in the wild-type strain. These findings suggest that the *recG* mutation plays a synergistic role with other genomic mutations to drive PCN increases. Taken together, this study presents the development of a pDNA hyperaccumulating *E. coli* strain with promising applications in industrial and therapeutic pDNA production, while also offering important insights into key genes involved in pDNA production.

## Introduction

As the field of gene therapy continues to advance, the demand for plasmid DNA (pDNA) has increased substantially, with industrial-scale applications requiring quantities ranging from hundreds of grams to several kilograms^1–5^. The low yield that can be isolated from *E. coli* strains, as well as the high cost of purification, have made pDNA production one of the main cost drivers for gene therapies, with prices reaching up to $100,000 per gram^4,6^. Significant efforts have been devoted to engineering the plasmid itself to increase pDNA production^7,8^. One example is the patented pNTC vector system, which achieves a remarkably high plasmid copy number (PCN) of approximately 1,200 copies per cell. This enhancement is enabled by the genomic integration of the toxic gene *sacB*, whose detrimental effects are suppressed by a weak RNA inhibitor encoded on the plasmid itself^7^. However, this vector-based modification approach requires a dedicated plasmid backbone and stable genomic insertion of a toxic gene, significantly limiting its flexibility and broader applicability. Given that plasmid replication relies on the host cell’s replication machinery and is tightly regulated within the cell, an alternative strategy is to engineer the native genes of *E. coli* production strains to boost plasmid synthesis^9^. This approach could work synergistically with existing vector technologies and offer a more versatile solution, enabling high-yield pDNA production from any user-supplied plasmid without necessitating modifications to the plasmid vector.

Plasmid replication is a complex process involving the participation of many host proteins^10^. To enhance plasmid production, researchers have pursued strategies aimed at improving plasmid stability within the host and increasing carbon flux toward nucleotide precursor synthesis^9,11^. Knockout *recA* and *endA* reduces plasmid degradation and unwanted recombination events^12^, while overexpression of *ligA* facilitates more efficient sealing of nicks during plasmid replication. Additionally, engineering central carbon metabolism genes, such as *pykA* and *pykF*, has been employed to boost flux through the pentose phosphate pathway and TCA cycle, thereby promoting nucleotide biosynthesis and reducing byproduct accumulation, such as acetate^9^. However, these studies mainly focus on engineering individual genes, overlooking potential interactions between genes that are involved in the process. Given the complexity of plasmid replication, systematically engineering strains by rationally engineering all combinations of potential genes would be infeasible. Instead, whole-genome random mutagenesis, combined with selection for beneficial variations, offers a more efficient approach^13–15^.

In this study, we employed a plasmid-based genome-wide mutagenesis approach to introduce random mutations across the genome and screened for variants with enhanced PCN. Specifically, we used a recently described anhydrotetracycline-inducible mutagenesis plasmid (aTc-MP6)^16^ to introduce mutations, followed by screening via a GFP reporter plasmid transformed to the mutated library. Variants with high cellular fluorescence, corresponding to high PCN, were recovered by fluorescence-activated cell sorting of over 10^6 variants. The best-performing strain, M3, exhibited increased PCN across all three tested origins (pUC, p15A, pSC101). Specifically, plasmids with pUC origins showed 5.93-fold (GFP reporter plasmid), 1.93-fold (gWiz DNA vaccine plasmid), and 8.7-fold (pAAV-CAGG-eGFP plasmid) increases. Plasmids with p15A and pSC101 origins exhibited 1.44-fold and 1.68-fold increases, respectively, compared with the parental NEB 5α strain. Among the 85 mutations identified in the adapted strain, MAGE (Multiplex Automated Genome Engineering) revealed the importance of *recG* (ATP-dependent DNA helicase) in plasmid production. Although the *recG* mutation alone did not enhance pDNA accumulation, its combination with other genetic alterations led to a significant increase in plasmid yield. These findings highlight the critical role of gene interactions in optimizing complex phenotypes and generate a promising strain for high-titer pDNA production.

## Materials and Methods

### Strains and media

Primers were obtained from Eurofins. NEB 10-beta was used for plasmid construction, while NEB 5α was the parental strain for genome mutagenesis. Bacterial strains were grown in Lysogeny Broth (LB) media (5 g/L yeast extract, 10 g/L tryptone, 10 g/l NaCl) at 37°C supplemented with ampicillin [(Amp) (100 µg/mL)] and kanamycin [(Kan) (50 µg/mL)] as appropriate. NEB 5α containing mutagenesis plasmid aTc-MP6 was grown in LB containing 1% (w/v) D-glucose and kanamycin. gWizPlum was a gift from Loree Heller (Addgene Plasmid # 74475), and pORTMAGE-3 was a gift from Csaba Pál (Addgene Plasmid # 72678)^17^. SPC822 and SPC1424 plasmids for CRISPR–Cas9–mediated genome editing were gifts from Scott Collins and Chase Beisel. pAAV-CAGG-eGFP plasmid was a gift from Edward Boyden (Addgene plasmid # 37825). All plasmids used in this study are listed in Table S1.

### Construction of a PCN reporter plasmid

The reporter plasmid was constructed using CIDAR MoClo Parts Kit (Addgene Kit #1000000059) as previously described^18^. Selected promoters and ribosomal binding sites were used for one-pot, multipart assembly to select the lowest nonzero GFP expression in NEB 5α. After transformation, 20 colonies were picked from the plate for fluorescence measurement using a BD Accuri C6 Plus Flow Cytometer. The plasmid from the colony exhibiting the lowest fluorescence was extracted using a Zyppy Plasmid Miniprep Kit and sent for full-length plasmid sequencing (https://www.plasmidsaurus.com/). To change the antibiotic resistance of the reporter plasmid, the resistance marker was switched from kanamycin (Kan) to ampicillin (Amp) using Gibson Assembly. Notably, even in widely used plasmid DNA production strains NEB 10β, a small fraction of plasmid multimers may be present. To prevent false positives during fluorescence-activated cell sorting (FACS), gel extraction was performed to isolate plasmid monomers before transformation.

### Induction of random mutations in the genome

NEB 5α was transformed with an anhydrotetracycline-inducible mutagenesis plasmid (aTc-MP6)^16^ and plated on LB agar containing 1% (w/v) D-glucose and kanamycin. A single colony from the plate was grown overnight in LB containing 1% (w/v) D-glucose and kanamycin and washed twice with PBS to remove glucose. Washed cells were transferred into new LB media (1:500 dilution) with kanamycin and 200 ng/mL aTc to induce mutagenesis. To investigate the number of mutations across different induction times, ten isolates were collected per five time points ( 2, 4, 6, 8, and 10 hours) post-induction and sent for whole-genome sequencing. After mutagenesis, cells were washed with PBS to remove aTc and resuspended in LB containing 1% (w/v) D-glucose and kanamycin. The mutated cell library was grown overnight to prepare competent cells for electroporation.

### High-efficiency electrotransformation of *E. coli*

Overnight cultures of mutated strains were inoculated into 100 mL LB broth (1:100) and grown to OD_600_ = 0.4 - 0.6. Cells were chilled on ice for 20 min and then harvested at 4000 x g for 15 min at 4 °C. Cell pellets were washed at least three times with 10 % ice-cold glycerol and resuspended in 1 mL 10 % glycerol. This suspension was divided into 50 μl aliquots and either put on ice for transformation immediately or stored at −80 °C. Frozen cells were thawed on ice for 5 min before the transformation. To achieve high transformation efficiency, 1 µg reporter plasmid was mixed with the competent cells and electroporated with a Biorad Micro Pulser electroporator at a time constant of approximately 5 ms. After 1 h recovery in SOC medium, transformed cells were either plated on LB with ampicillin plates to count CFU/mL or subcultured (1:50 dilution) in LB media with ampicillin for cell sorting.

### Fluorescence-activated cell sorting (FACS)

Flow cytometric analysis and cell sorting were performed using a BD FACS Melody. Cells were harvested during the mid-log phase, washed twice with PBS, and then resuspended in PBS. They were kept on ice prior to sorting. NEB 5α transformed with reporter plasmid was used to set the gates, voltages, and thresholds. 1000 cells were sorted and plated across ten LB agar plates with Amp (around 100 colonies on each plate). Each plate was visualized under a UV flashlight, and colonies with higher fluorescence were picked from the plate for further analysis.

### Whole genome sequencing and PCN calculations

Overnight cultures of both parental NEB 5α and sorted variants with reporter plasmid were inoculated in 25 mL LB media with Amp and grown for 15 h. Saturated cultures were subinoculated in 25 mL LB media (Amp) with an initial OD_600_ = 0.01 and grown for 8 h. 1 mL cell culture was spun down at 3000 x g for 5 mins, and cell pellets were resuspended in 200 µl nuclease-free water for gDNA extraction with Zymo Quick-DNA Fungal/Bacterial Miniprep Kit. gDNA samples were sent to SeqCenter for Illumina Whole Genome Sequencing, generating 200 Mbp 2×151bp paired-end reads per isolate. Sequencing reads were aligned to the NEB 5α reference genome (GenBank: CP017100.1) and the corresponding plasmid sequence within the cell. Mutation detection was performed using breseq with default settings. To estimate PCN, we used the “fit mean” metric generated by breseq^19^, which reflects the average sequencing coverage depth across the genome. The relative PCN was determined by calculating the ratio of average plasmid coverage to chromosomal coverage.

### Agarose gel electrophoresis for analyzing plasmid topology

The presence of the supercoiled circular (SC) plasmid on a gel was determined by preparing pDNA to the open circular (OC) form through nicking or to the linear form using restriction enzyme digestion. To nick the dsDNA to the OC form, 1 µL of NtBstNBI was combined with 1 µg of dsDNA and 5 µL of 10X NEBuffer r3.1. NF water was added to reach a total volume of 50 µl, followed by incubation at 55 °C for 1 hour. The procedure was similar for obtaining linearized plasmid, except that 1 µL of BsaI-HFv2 was added instead of NtBstNBI, and the mixture was incubated at 37 °C for 15 min. Agarose gel electrophoresis was conducted to assess plasmid topology. A 1% agarose gel was prepared by dissolving 0.3 g agarose powder in 30 mL 1X TAE buffer and then poured into a gel casting tray with a comb for well formation. DNA samples, mixed with 6X loading dye, were loaded into the wells and electrophoresed at 100 V for 30 min before visualization.

### CRISPR-Cas9-mediated deletion of *recA* from NEB 5α

The *recA* gene was deleted from the NEB 5α genome using CRISPR-Cas9-mediated genome editing, following previously described methods^20^. An overnight culture of NEB 5α transformed with SPC822 plasmid (carrying Cas9 and λ Red recombination system under an IPTG-inducible lac promoter, a pBAD promoter driving expression of a gRNA targeting the ColE1 origin for curing SPC1424, and a temperature-sensitive pSC101 origin) was subcultured into 50 mL LB medium supplemented with streptomycin and 1 mM IPTG to induce Lambda RED expression. Electrocompetent cells were prepared, and 1 μg of the repair template and 100 ng of SPC1424 plasmid (expressing sgRNA targeting *recA*) were electroporated into the cells. Following electroporation, the cells were recovered in SOC medium at 30 °C for 4 hours before plating on LB agar for 24 hours. Successful *recA* deletion was confirmed by colony PCR. To cure SPC1424, a sequence-verified colony was cultured in 1 mL LB supplemented with 0.2% (w/v) arabinose to induce the expression of gRNA targeting the ColE1 origin. The SPC822 plasmid was subsequently removed by culturing the strain at 42 °C, utilizing its temperature-sensitive origin. The repair template used for *recA* deletion was generated by overlap PCR^21^. The primers for overlap PCR and sgRNA sequences are listed in Table S2.

### Bioreactor cultivation

Batch fermentation was carried out using Sartorius Biostat® B bioreactors at the Golden LEAF Biomanufacturing Training and Education Center (BTEC) at NCSU. The fermentation media was LB media with Amp, and the working volume was 1 L with the initial OD_600_ = 0.05. 0.1% antifoam 204 was added to avoid foam generation, and dissolved oxygen (DO) probes were calibrated before inoculation to 100% saturation at 400 rpm. The fermentation was controlled to 37 °C, aeration at 1 slpm, and dissolved oxygen at 30% with agitation recruited in the DO control cascade. 1 mL of cell culture was taken from the bioreactor every 1 h for plasmid extraction. All fermentations were run in triplicate.

### MAGE experiments

In order to identify the mutations responsible for the increased PCN, 10 consecutive MAGE cycles were conducted using mixed homologous oligos targeting all 85 sites. For each MAGE cycle, NEB 5α Δ*recA* harboring pORTMAGE-3^17^ was grown at 30 °C in a shaking incubator at 250 rpm until OD_600_ reached 0.5–0.7. Then, the culture was shifted to 42 °C and incubated for 15 min to induce expression of λ-Red recombination proteins. Cells were then immediately chilled on ice for at least 20 min and washed three times with 10% glycerol. 50 μL of cell suspension was mixed with oligos in single-plex experiments. Immediately after electroporation, 1 mL SOC medium was added for recovery. After 60 min at 30 °C and 250 rpm, 5 mL LB media was added and cells were allowed to reach mid-logarithmic state under continuous agitation, and this concluded one MAGE cycle. After 10 consecutive MAGE cycles, cells were transformed with the reporter plasmid before sorting. To evaluate the off-target mutation rate of MAGE, 10 random isolates were selected before the transformation with the reporter plasmid.

## Results and Discussion

### Overview of inducible genome-wide mutagenesis for pDNA accumulation

During normal DNA replication in *E. coli*, mutation rate is controlled from 10^−9^ to 10^−11^ errors per base pair^22^. *In vivo* mutagenesis methods increase this mutation rate and accelerate laboratory evolution, eliminating the need for labor-intensive *in vitro* manipulations^23^. Unlike chemical mutagenesis methods like N-methyl-N’-nitro-N-nitrosoguanidine and ethyl methanesulfonate, or physical methods such as ultraviolet light, which have limitations due to cellular toxicity and narrow mutagenesis spectrum, plasmid-based methods offer a tunable mutation rate, lower toxicity, and a broader mutational spectrum^16,23^. Of these, the widely used mutagenesis plasmid MP6 has been applied to evolve antibiotic resistance^23^, enhance fitness in deuterium-based growth media^15^, and improve biosorption of rare earth elements (REE)^24^. Building on the advantages of plasmid-based mutagenesis systems, we aimed to enhance plasmid DNA (pDNA) yield by introducing the aTc-inducible MP6 (aTc-MP6)^16^ into *E. coli* NEB 5α, a widely used host for pDNA production. Upon induction with anhydrotetracycline (aTc), this system enables the generation of random mutations *in vivo*, potentially facilitating adaptive improvements in pDNA titer. High pDNA-producing variants could then be identified using a plasmid-based reporter system that reflects PCN (Fig. 1A).

**Figure 1:**
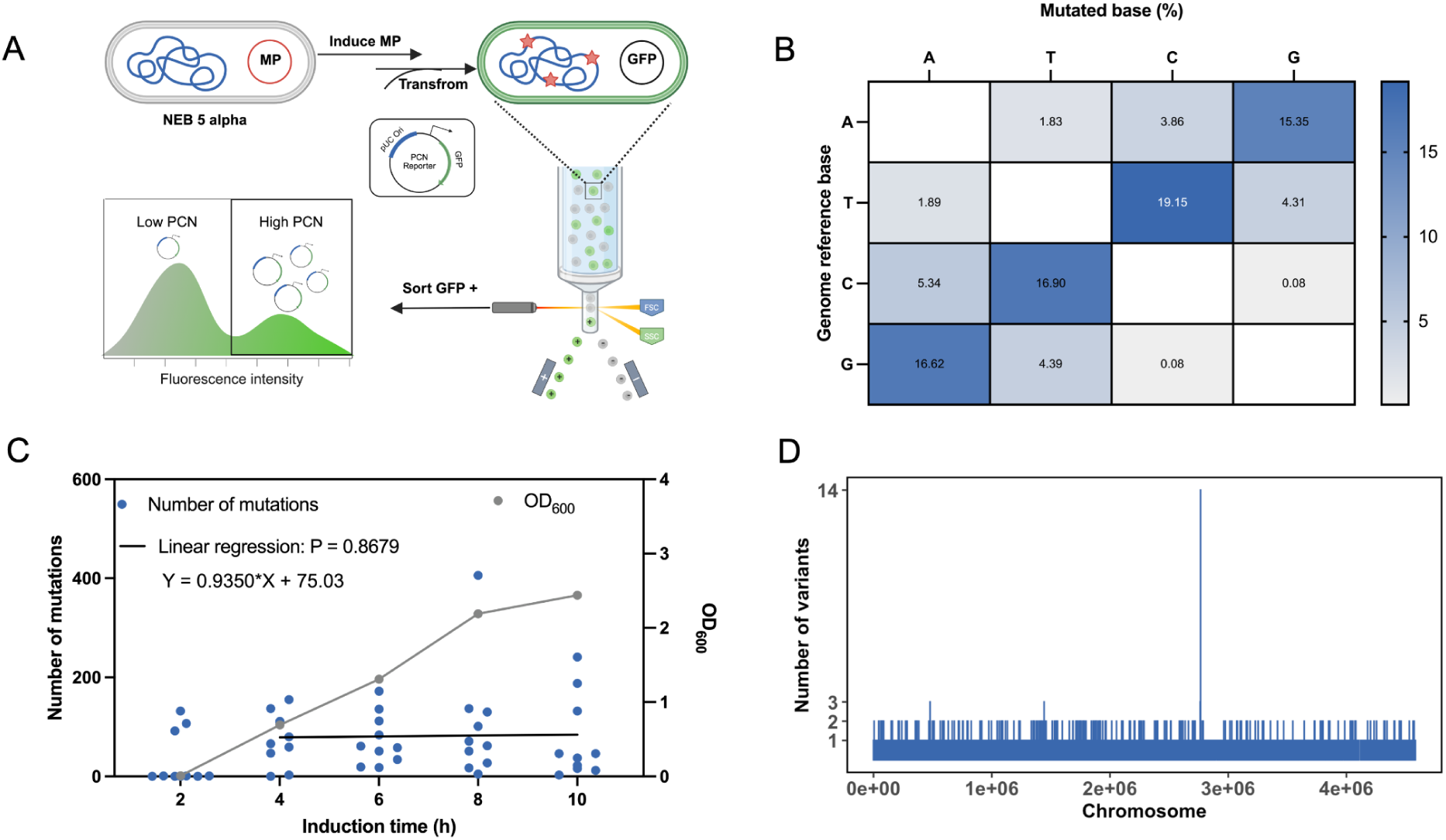
(A) Overview of inducible genome-wide mutagenesis. Random mutations were introduced into NEB 5α upon aTc induction. Strains with increased PCN were isolated using FACS. Figure made with BioRender (https://www.biorender.com/). (B) The percentages of base pair mutations between different nucleotides (A, T, C, G). Each cell represents the percentage of mutations from the nucleotide in the corresponding row to the nucleotide in the corresponding column. (C) The number of mutations observed across ten isolates recovered from different induction times (2, 4, 6, 8, 10 hours). (D) Distribution of gene mutations across chromosomes, with the x-axis representing the chromosomal position and the y-axis indicating the number of mutations.

### Characterization of genomic mutation spectra

Before applying this genomic mutagenesis method for the improvement of pDNA production, we analyzed the types of mutations introduced by aTc-MP6 within the genome. Previous studies of mutagenesis plasmids have primarily focused on mutations in *rpoB* using the rifampin-resistance assay, which offers simplicity but lacks a comprehensive view of genomic alterations^15,16,23,25^. To identify where the exact mutations are and which substitutions are introduced, we conducted whole-genome sequencing on 50 randomly selected isolates after 2 h, 4 h, 6 h, 8 h, and 10 h of induction to identify the mutation sites and mutagenic spectra. Base pair substitutions constituted 90% of all mutation types (Fig. 1B), with the remainder being insertions and deletions (indels) occurring within homopolymeric sequences exceeding four consecutive identical nucleotides. Unlike the mutation spectrum observed for MP6 via *rpoB*^23^, which showed 9 base change mutations with a balanced distribution of transitions and transversions, aTc-MP6 induces all 12 possible base change mutations across the genome, and displays a strong bias toward transitions between T:A↔C:G base pairs. This mutation pattern is likely driven by the impairment of mismatch repair mechanisms via the overexpression of two genes encoded on MP6: *dam* and *seqA*^23,26^.

To investigate the onset of mutation accumulation, cells were induced for varying durations (2, 4, 6, 8, and 10 hours) (Fig. 1C). Within the initial 2-hour induction period, most cells had not undergone replication, and consequently, no genetic mutations occurred. However, in some isolates (3 out of 10), approximately 100 mutations were detected. These isolates could represent cells that were in the initial stages of replication and mutation accumulation. As the induction time increased to 4 hours, a significant accumulation of mutations occurred, with many more isolates containing mutations. Further extension of the induction time did not lead to a further increase in mutations (Figure 1C). This phenomenon may be attributed to cells reaching the stationary phase and halting replication, subsequently limiting additional mutations, or it could be due to reaching a maximum tolerable number of mutations before substantial reductions in cell viability occur.

Notably, we found that aTc-MP6 induces mutations throughout the entire chromosome (Fig. 2D). However, among 50 randomly selected isolates, 14 shared the identical *srlA* F168L mutation. This enrichment may result from *srlA* being part of a network of genes involved in stress-induced mutagenesis^27^. Under stress conditions, genes involved in stress responses may become more transcriptionally active^28,29^, potentially increasing their susceptibility to MP6-induced mutagenesis. Although the F168L mutation in *srlA* does not significantly impact the overall mutation rate induced by MP6 (Fig. S1A), it may act as a pleiotropic regulator, influencing cell growth during mutation induction and conferring a selective advantage in these settings.

**Figure 2:**
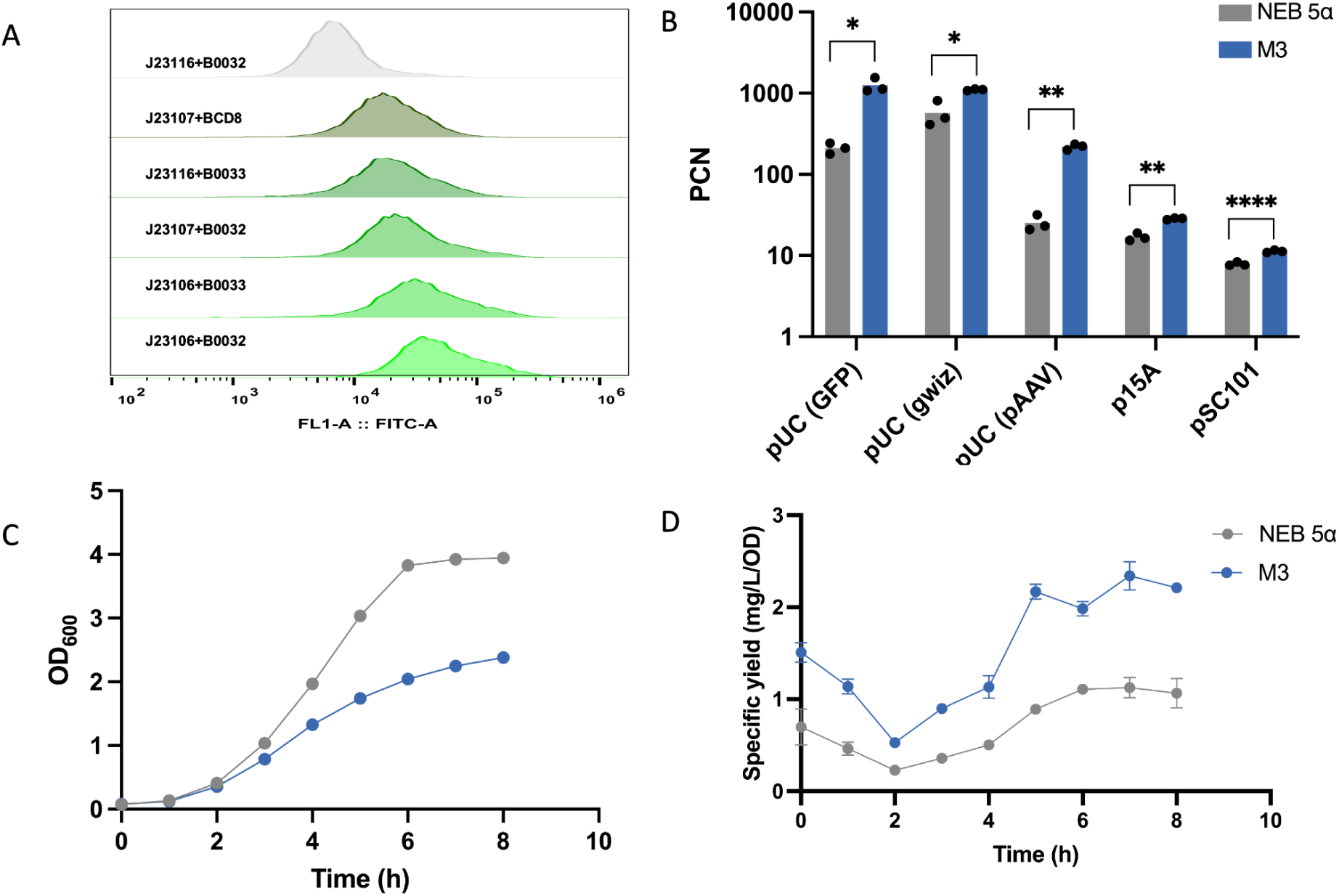
(A) Flow cytometry histograms of candidate PCN-responsive screening plasmids. The J23116 promoter with the B0032 RBS showed the weakest fluorescence within the cell. (B) The PCN of NEB 5α and M3 for three pUC-based plasmids (GFP reporter plasmid, gWiz DNA vaccine plasmid, and pAAV-CAGG-eGFP plasmid), and plasmids with the p15A origin and pSC101 origin. (C) Over the course of 8 hours, M3 demonstrated a reduced growth rate and attained a lower final cell density compared to NEB 5α in 1L bioreactor fermentations. (D) M3 exhibited a higher specific yield throughout the fermentation process, with a 2.3-fold increase in specific yield at 8 hours versus the parental strain.

### Construction of a PCN reporter plasmid

To select variants with increased PCN, we constructed a GFP reporter plasmid containing the pUC origin of replication, a commonly used origin for DNA vaccine and gene therapy production. Due to the high native PCN of pUC-based plasmids within the cell, keeping GFP expression low for the parental strain was essential. Failing to do so could lead to difficulties in identifying variants, as GFP expression would already be saturated before mutagenesis. To obtain a GFP plasmid with weak fluorescence, we assembled combinations of three weak promoters (J23106, J23107, and J23116) and three weak ribosomal binding sites (B0032m, B0033m, and BCD8) from the CIDAR MoClo Parts Kit^18^. Of 20 colonies recovered after transformation (Fig. S1B), we assessed six distinct combinations of promoters and RBSs (Fig. 2A). Among these, we found that the combination of promoter J23116 and RBS B0032 exhibited the lowest expression levels and, therefore, this combination was selected for GFP expression.

### *recA1* reversion leads to plasmid multimerization

With the reporter plasmid ready, NEB 5α cells harboring aTc-MP6 were induced using aTc and then transformed with the reporter plasmid to isolate highly fluorescent cells. Two variants (M1 and M2) isolated from FACS exhibited enhanced fluorescence levels. Upon curing aTc-MP6, the reporter plasmid was extracted from both mutants and wild-type NEB 5α and loaded onto an agarose gel to check the plasmid topology (Fig. S1C). Compared with NEB 5α, which mainly produces supercoiled monomers (∼ 3 kb), both variants produced plasmid multimers (M1: dimers and tetramers; M2: trimers, pentamers, and hexamers). This gel result was consistent with the whole plasmid sequencing results, which also showed the formation of plasmid multimers during long-read sequencing (Fig. S2A-C).

To identify the genes responsible for multimerization, the genomic DNA of the variants was subjected to whole-genome sequencing. Typically, NEB 5α possesses a *recA1* (G161D) mutation that decreases recombination^30^. However, in the variants that produced multimers, the *recA1* mutation had reverted (D161G) and therefore likely increased recombination as a strategy for increasing per-cell fluorescence.

### Inducible genome evolution of NEB 5α *ΔrecA* yields a strain harboring high-copy monomeric plasmids

We hypothesized that completely removing the *recA* gene would prevent the reversal of the *recA1* mutation during mutagenesis and would allow for the selection of a variant with a higher PCN but without multimerization. To test this, the entire *recA* gene was deleted from NEB 5α using CRISPR-Cas9-based genome editing^20^, and the resulting strain (NEB 5α *ΔrecA*) was used as the starting strain for a new round of evolution. Using this approach, a new variant (M3) with increased GFP expression was obtained after FACS. As expected, the extracted reporter plasmid revealed the same supercoiled monomeric topology as NEB 5α (Fig. S2D).

We assayed the PCN enabled by M3 using next-generation sequencing using three different plasmids harboring pUC origins: a GFP reporter plasmid, the gWiz plasmid^31^ designed for mammalian expression, and pAAV-CAGG-eGFP^32^, an adeno-associated virus (AAV) vector for gene therapy applications. As expected, M3 enabled increased PCN across all three constructs compared to NEB 5α (Fig. 2B). To determine whether this enhancement extended to plasmids with alternative replication origins, we replaced the pUC origin in the reporter plasmid with either p15A or pSC101. Interestingly, M3 also promoted elevated PCN for both p15A and pSC101 origins. This broader enhancement may result from M3’s modification of host factors that are shared among multiple replication systems, such as increased availability of replication initiation proteins, reduced plasmid replication control, or improved plasmid stability.

The performance of NEB 5α and M3 was next evaluated in small-scale 1 L bioreactors to measure pDNA titers at larger culture volumes. Over a short (8 h) cultivation, M3 exhibited a slower growth rate compared to NEB 5α and achieved a lower final density (Fig. 2C). This could be attributed to the increased metabolic burden caused by increased PCN^33,34^. However, M3 demonstrated a higher specific yield during the fermentation process (2.3-fold increase for specific yield at 8 h, Fig. 2D). The fold change in specific pDNA yield in the bioreactor (2.3-fold) was lower than the fold change in PCN observed in the shake flask (5.93-fold), which may be due to the inefficiency of pDNA purification and/or differences between shake flask and bioreactor fermentation conditions^4,35^. Further optimization efforts (e.g., culture time, media composition, temperature, aeration, pH) are crucial to enhance pDNA yield^36–39^.

### Multiplex Automated Genome Engineering (MAGE) reveals a *recG* mutation as a key determinant of high PCN

To elucidate the genetic basis for the increased PCN in strain M3, we performed whole-genome sequencing and identified 85 mutations in the genome (Table S3). To pinpoint the mutations responsible for this phenotype, we employed multiplex automated genome engineering (MAGE) to systematically reconstruct combinatorial variants of these mutations in the NEB 5α background^17,40–43^. After conducting 10 cycles of MAGE, we identified 7 isolates (M4, M5, M6, M7, M8, M9, M10) with increased PCN relative to the NEB 5α parental strain, although none reached the levels observed in M3 (Fig. 3). This suggested that 10 cycles may not have been sufficient to explore all relevant combinations of mutations, and additional cycles may be necessary to fully reconstruct the genetic determinants of the M3 phenotype. Whole-genome sequencing revealed that all seven isolates harbored a common *recG* R584H mutation. To assess the contribution of this mutation, we introduced it into NEB 5α (yielding strain M11), which did not lead to increased plasmid production. However, reverting the *recG* R584H mutation in M3 resulted in a 63% reduction in plasmid yield, indicating that this mutation alone is insufficient but acts synergistically with other genetic changes present in M3 to enhance plasmid DNA (pDNA) replication.

**Figure 3:**
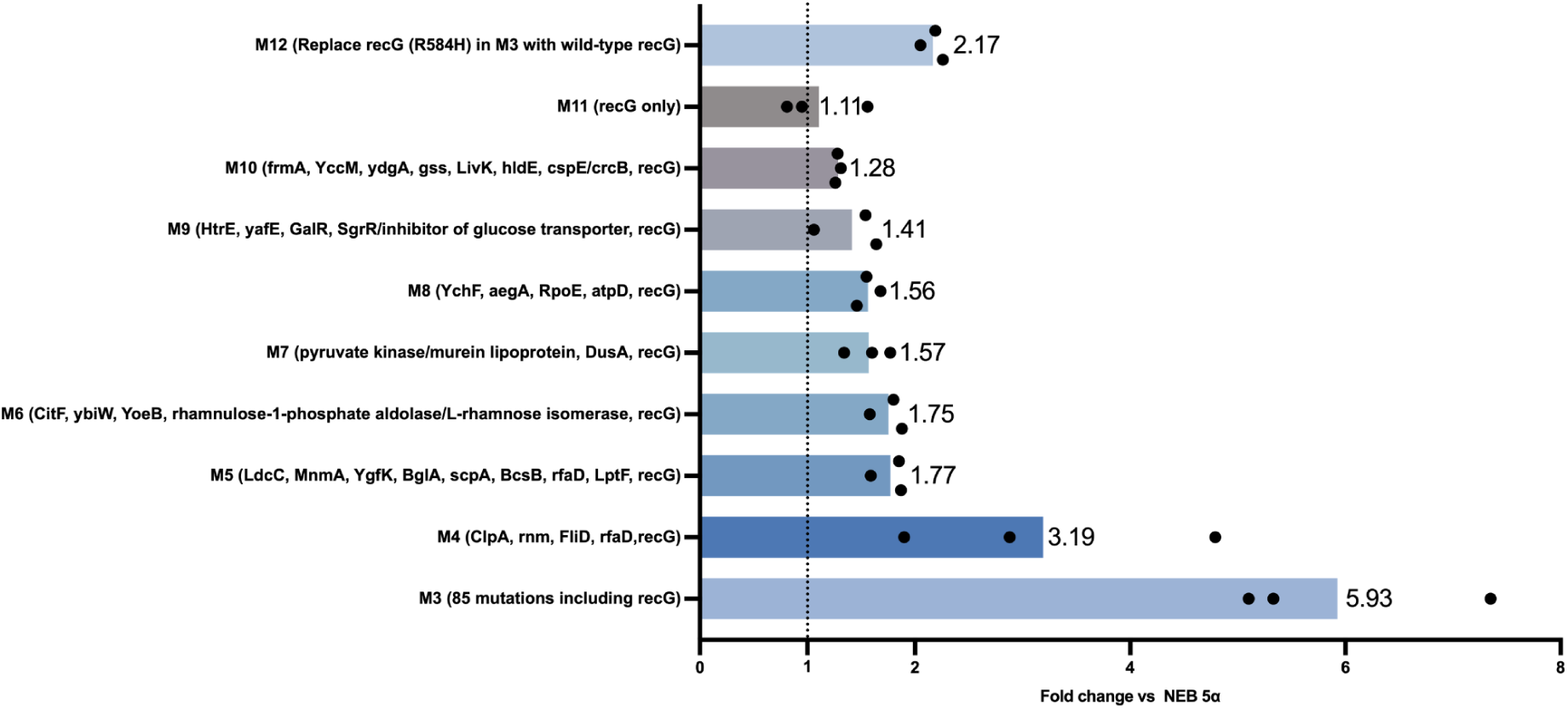
Relative PCN fold change of M3–M12 compared to NEB 5α. M3 shows the highest increase in PCN, whereas M11 (*recG* R584H alone) does not differ significantly from NEB 5α. Reversion of *recG* R584H in M3 (M12) led to a marked reduction in PCN, highlighting the context-dependent role of *recG* in plasmid replication.

*recG* encodes a helicase involved in DNA repair and replication fork regression^44^. The R584H substitution may alter RecG function in a way that, in the context of additional mutations, promotes a cellular environment favorable to plasmid replication—potentially by modifying replication fork dynamics, relieving stress responses, or indirectly enhancing plasmid-specific replication mechanisms. These findings suggest that the effect of *recG* on plasmid yield is shaped by epistatic interactions within the evolved genomic background of M3.

## Conclusion and discussion

A plasmid-based mutagenesis strategy was evaluated for inducible genome-wide mutagenesis. The broad mutational spectrum of aTc-MP6, along with its uniform genomic distribution, enabled random mutagenesis across the genome within 4 hours. This approach can also be used for accelerating adaptive laboratory evolution, a process that typically requires weeks to months to accumulate mutations. When applied to enhance the PCN of plasmids with pUC origins in *E. coli* strains, we initially observed a reversion of the *recA1* mutation, which led to plasmid multimerization. However, for DNA vaccine production, a supercoiled plasmid monomer is preferred. Thus, complete *recA* knockout was necessary for further strain optimization, as the *recA1* mutation involves only a single base pair change and is therefore easy to revert.

Our experiments indicated that a single nucleotide change within *recG* was key to improving PCN. Previous studies of *recG* have primarily focused on gene deletion—often in combination with a second gene of interest—to assess phenotypic consequences^45,46^. However, this approach overlooks the impact of single-nucleotide mutations, which occur more frequently during natural evolution, particularly in essential genes where complete deletions are often lethal or highly deleterious. Moreover, such deletion-based studies provided only a narrow and fragmented view of the genetic interactions involving *recG*. In our work, we identified a total of 84 potential mutations that may enhance PCN in the context of *recG* variation, highlighting the complex regulatory network governing plasmid replication. Given that our parental strain, NEB 5α *ΔrecA*, was already engineered with beneficial mutations for plasmid production, it is unsurprising that extensive mutational changes were required to achieve further enhancement. The successful recovery of this high-performing strain validates both the effectiveness of this screening strategy and the potential of genome-wide mutagenesis for identifying novel engineering targets. The engineered strain demonstrated improved compatibility and performance across three different plasmid origins—pUC, p15A, and pSC101—indicating its broad utility. Although the strain’s performance during our bioreactor tests (2.3 fold increase versus wild-type) was less than anticipated, we should note that commercial pDNA fermentation often occurs over a much longer time (8 h batch vs over 30 h fed batch) and process parameters, including feeding rates and media composition, are usually tailored in a strain-specific manner. Thus, we expect that after culture optimization, this strain will be suitable not only for DNA vaccine production but also for high-yield plasmid preparation, particularly for plasmids with inherently low PCN, with potential applications in both laboratory and industrial settings.

Although Multiplex Automated Genome Engineering (MAGE) is a powerful strategy for studying essential genes, testing all possible combinations of mutations across the 85 candidate genes we identified was not feasible (2^85 = 3.9*10^25), limiting our search to smaller subsets of the 85 mutations. Additionally, despite using an inducible mismatch repair (MMR) inactivation system to reduce editing bias, off-target mutations were still observed after 10 cycles of MAGE. Indeed, in a sample of 10 random isolates picked (Figure S3), the average number of on-target mutations was 2.9, whereas the average number of off-target mutations was 4.9. Increasing the number of MAGE cycles may further elevate off-target mutation rates, complicating the interpretation of causative variants.

Taken together, we characterized a genomic mutagenesis approach and successfully used it to engineer a plasmid hyperproduction strain. This strain harbors 5.93-fold more pUC-based GFP reporter plasmid within its cell than its parental strain, but grows significantly slower. Additional optimization of culture conditions may enable enhanced growth of this strain and concomitant increases in volumetric plasmid titers. Analysis of the resulting mutations in this strain revealed a complex genetic network involving *recG*. While current limitations in the number of sequenced *E. coli* mutants prevent a full dissection of this intricate interaction landscape in the context of pDNA production, future technological advancements may enable a more comprehensive understanding of this *recG*-centered regulatory network governing plasmid replication.

## Acknowledgments

We thank Drs. Ryan Paerl and John van Schaik for FACS melody training and Haiwei Zhang and Melody Lao from BTEC for carrying out the fermentation. We also thank the members of the Crook Lab for helpful discussions. This work was supported by the North Carolina Biotechnology Center (2022-TRG-6707).

## Supplementary

**Figure S1:**
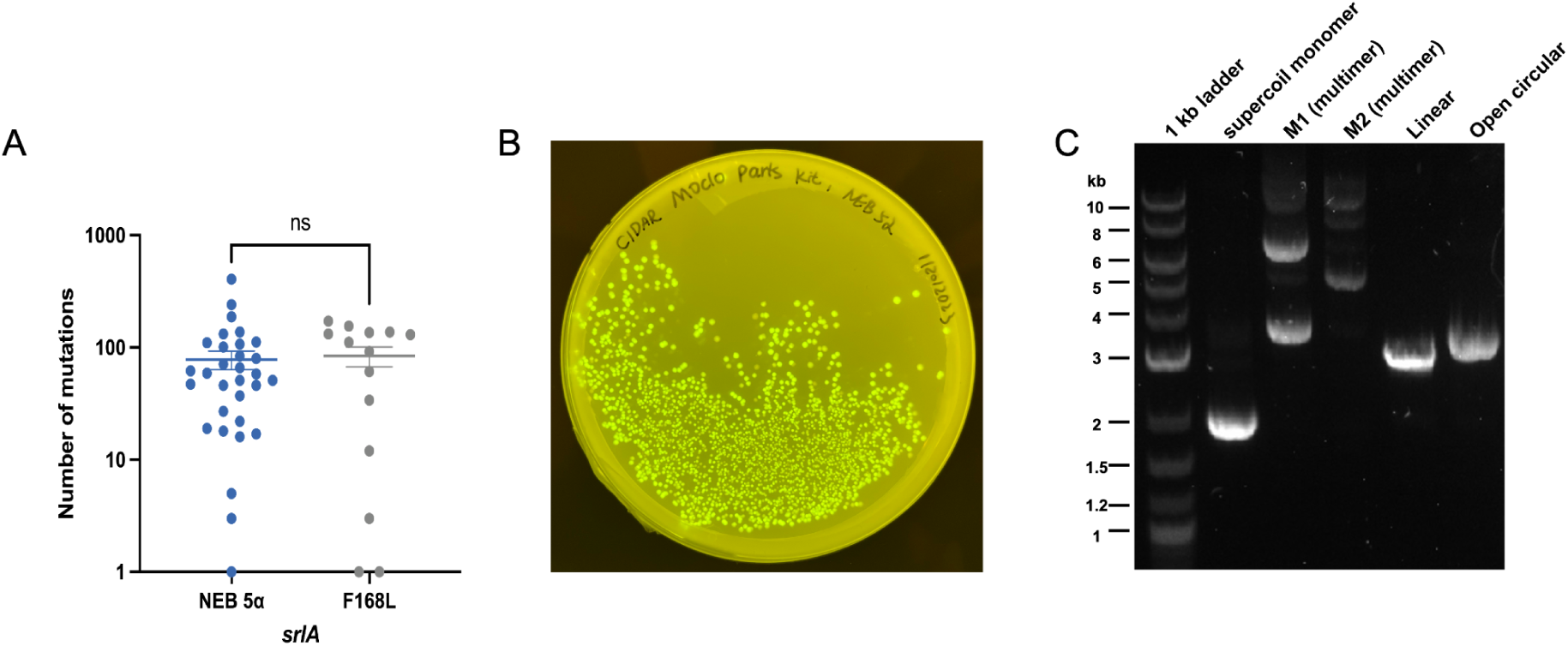
(A) The F168L mutation in srlA does not significantly affect the overall mutation rate induced by MP6. The number of mutations induced by MP6 was quantified in WT (without the srlA F168L mutation) and the srlA F168L mutant. Each dot represents an individual isolate randomly selected (n = 50) after induction, with isolates containing zero mutations excluded. Data are presented as mean ± standard error of the mean (SEM). Statistical analysis shows no significant difference between WT and srlA F168L (P > 0.05, ns). (B) Fluorescence intensity of colonies after transformation, showcasing varying brightness levels due to the distinct combinations of promoters and RBS. (C) Gel electrophoresis image of reporter plasmids. Line 1: 1 kb DNA ladder. Line 2: Plasmid extracted from NEB 5α, primarily producing supercoiled monomers. Line 3: Plasmid extracted from M1. Line 4: Plasmid extracted from M2. Line 5: Linearized plasmid. Line 6: Open circular plasmid.

**Figure S2:**
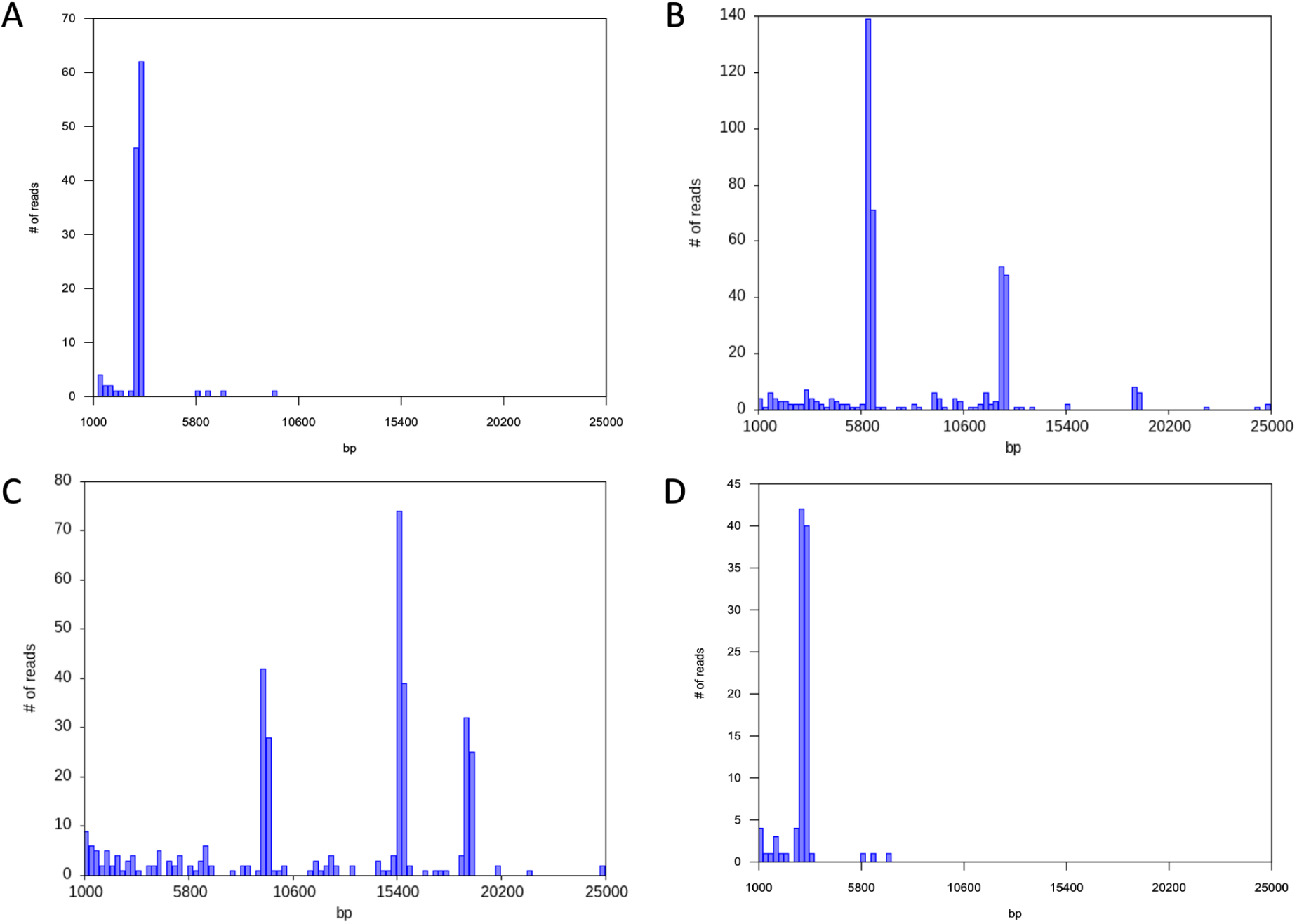
Read length histograms of the reporter plasmid extracted from NEB 5α, M1, and M2. (A) Plasmid extracted from NEB 5 alpha shows a dominant peak at 3.1 kb, corresponding to the plasmid monomer. (B) Plasmid extracted from M1 exhibits two peaks, indicating the presence of dimers and tetramers. (C) Plasmid extracted from M2 displays three peaks, corresponding to trimers, pentamers, and hexamers. (D) Plasmid extracted from M3.The single peak, similar to the parental NEB 5α strain, indicates the presence of the plasmid monomer.

**Figure S3:**
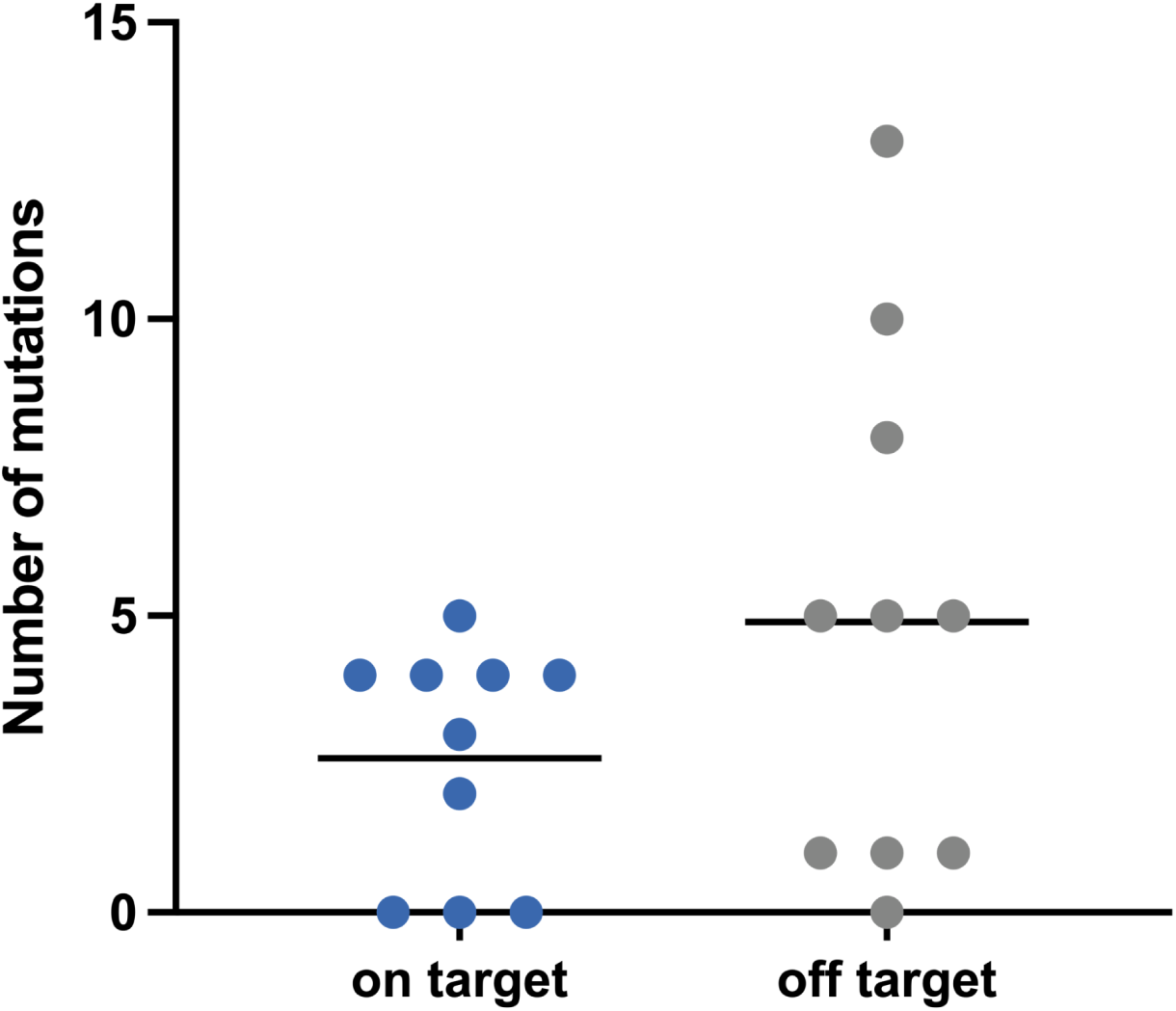
Analysis of on-target and off-target mutations after 10 cycles of MAGE. 10 random isolates were picked prior to reporter plasmid transformation to evaluate mutation outcomes. On average, each isolate carried 2.9 on-target mutations and 4.9 off-target mutations, indicating a higher frequency of off-target events after multiple MAGE cycles.

